# *Bartonella* and hemotropic *Mycoplasma* species in synanthropic bats in Kenya

**DOI:** 10.64898/2026.05.28.728508

**Authors:** Isabella K. DeAnglis, Tamika J. Lunn, Reilly T. Jackson, Caroline A. Cummings, Emmarie C. Gates, Benjamin Griffey, Benson Mwakachola, Peter Mwasi, Joseph G. Ogola, Paul W. Webala, Tarja Sironen, Daniel J. Becker, Kristian M. Forbes

## Abstract

Identifying and characterizing zoonotic pathogens in wildlife is essential for understanding disease risk to humans. In Sub-Saharan Africa, many people live with bats in their houses and are exposed to their pathogens, yet little is known about the bacterial pathogens in Afrotropical bat species. Globally, *Bartonella* spp. (bartonellae) and hemotropic *Mycoplasma* spp. (hemoplasmas) are common bacterial pathogens in bats, and some lineages are known to spill over and cause infections in humans. To evaluate this disease risk, we screened three common synanthropic bat species in Kenya, and their ectoparasites, for hemoplasmas and bartonellae and assessed their relatedness to known human pathogens. Of 767 bats across 21 sites, 17.9% of bats were *Bartonella* spp. positive and 19.3% were hemoplasma positive. Bat ectoparasites had similar *Bartonella* prevalence (13.5–25.0%) and, for most bat species, ectoparasite loads were not associated with increased likelihood of *Bartonella* infection. We found that *Bartonella* lineages displayed phylogenetic overlap between different bat species and ectoparasites, suggesting pathogen sharing between species, while hemoplasma lineages corresponded strictly to host taxonomy. Finally, we found that 16S rRNA sequences from one heart-nosed bat (*Cardioderma cor*) were 97.85% similar to a human-associated hemoplasma found previously in Schreiber’s bats (*Miniopterus schreibersii*) in Spain. We show that synanthropic bats host bacteria of potential public health concern, highlighting the need to investigate the emerging impacts of these pathogens on human health in Kenya and elsewhere in Sub-Saharan Africa.

## Introduction

Identifying and characterizing zoonotic pathogens in wildlife is essential for predicting disease risk to humans. In urbanizing areas, habitat loss and fragmentation alter the spatial distribution of wildlife populations (Buchmann *et al*. 2013). These processes reduce suitable habitat for wildlife and often increase human–wildlife contact (Hassell *et al*. 2017). Synanthropic wildlife—species that have adapted to living in human-dominated landscapes—are often found in and around anthropogenic buildings, resulting in direct and indirect contact between wildlife and humans (Voigt *et al*. 2015; Gravinatti *et al*. 2020; Jackson *et al*. 2024). Because of these frequent contacts, humans are regularly exposed to synanthropic wildlife and their pathogens, increasing the likelihood of zoonotic spillover (McFarlane *et al*. 2012; Ecke *et al*.2022; Fraint *et al*. 2025). Knowing what pathogens are circulating in synanthropic wildlife, and whether these pathogens are infectious to humans, is essential to developing public health strategies that protect people from potential outbreaks.

Globally, bats are among the most common and widespread synanthropic wildlife, often seeking out human structures for stable roosting habitat (Entwistle *et al*. 1997; Betke *et al*. 2024). Bats also host of a wide diversity of pathogens, including pathogens documented to cause disease in humans (Mühldorfer *et al*. 2013; Chomel *et al*. 2014; Allocati *et al*. 2016). However, to date, most bat-borne disease research—a topic that has seen a large increase in research effort over the past decade—focuses on viral pathogens (Letko *et al*. 2020). Bacterial pathogens have been comparatively neglected, despite a diversity of known bat-borne bacteria with zoonotic potential that remain poorly characterized and understood (Mühldorfer *et al*. 2013; Szentivanyi *et al*. 2023).

In terms of zoonotic risk, *Bartonella* spp. and hemotropic *Mycoplasma* spp. (hereafter hemoplasmas) represent two of the most consequential bat-borne bacterial taxa, as they are often highly prevalent within bat populations in some geographic regions (Becker *et al*. 2018; Szentivanyi *et al*. 2023; Wang *et al*. 2023) and cause erythrocytic infections in a variety of mammals, including humans (Messick 2008; Lin *et al*. 2010; Descloux *et al*. 2021). Bat-borne hemoplasmas, such as *Candidatus* Mycoplasma haematohominis and a *Mycoplasma* sp. from a Schreiber’s bats (*Miniopterus schreibersii*) have been detected in humans in Asia, Oceania, and Europe (Millan *et al*. 2015; Hattori *et al*. 2020; Descloux *et al*. 2021; Esperon *et al*. 2024), and bat-borne *Bartonella* spp., such as *Candidatus* Bartonella mayotimonensis and *Candidatus* Bartonella rousetti, have been detected in humans in North America, Europe, and Africa (Lin *et al*. 2010; Veikkolainen *et al*. 2014; Bai *et al*. 2018), indicating zoonotic transmission. *Bartonella rochalimae*, a known human pathogen (Eremeeva *et al*. 2007), was also recently identified in North American bats (Becker *et al*. 2025). Arthropod ectoparasites, including fleas, mites, ticks, and bat flies, have been implicated in the transmission of these bacteria within animal populations (Willi *et al*. 2007; Millan *et al*. 2021; McKee *et al*. 2026), but the specific role of vectors in maintenance and transmission of these bacteria remains largely unclear. Although these bacteria can cause anemia, fever, and significant morbidity in immunocompromised people, little is known about the ecological drivers of their maintenance and transmission (Szentivanyi *et al*. 2023). Furthermore, few studies have investigated these bacteria in Africa (Kamani *et al*. 2014; Brook *et al*. 2015; Dietrich *et al*. 2016; Bai *et al*. 2018) and rarely have they been investigated in synanthropic bats (but see Kosoy *et al*. 2010; Di Cataldo *et al*. 2020) that have regular contact with humans.

Afrotropical bats make up approximately 20% of the world’s known bat species, yet research on their bacterial pathogens remains limited, especially in insectivorous species (Szentivanyi *et al*. 2023; Monadjem *et al*. 2024). Some studies have begun to reveal diverse bat-associated bartonellae in South Africa, Madagascar, Nigeria, and Kenya (Kosoy *et al*. 2010; Kamani *et al*. 2014; Brook *et al*. 2015; Dietrich *et al*. 2016; Bai *et al*. 2018), suggesting that these bacteria could be distributed across the continent. Additionally, one study in Nigeria found hemoplasmas are prevalent, widely distributed, and genetically diverse in bats (Di Cataldo *et al*. 2020), but very little is known about bat-borne hemoplasmas in other African countries. This limited previous work on these bacterial pathogens in Africa has also focused on cave- and forest-dwelling populations of bats, despite the limited human–wildlife contact that occurs with these species compared to synanthropic species (Kamani *et al*. 2014; Brook *et al*. 2015; Dietrich *et al*. 2016; Bai *et al*. 2018). Furthermore, African studies investigating the vectors of these bat-borne bacteria have also been restricted and mainly focus on bat flies that parasitize fruit bats, so little is known about other ectoparasites that may vector these bacteria in anthropogenic settings (Kamani *et al*. 2014; Brook *et al*. 2015; Dietrich *et al*. 2016).

To characterize bat-borne bacterial pathogens and investigate the risk they pose to humans in Sub-Saharan Africa, we screened three common synanthropic bat species, Angolan free-tailed bats (*Mops condylurus*), little free-tailed bats (*Mops pumilus*), and heart-nosed bats (*Cardioderma cor*) in rural Kenya, as well as their ectoparasites, for bartonellae and hemoplasmas. We conducted phylogenetic analyses to assess the relatedness of the bacteria we found to other lineages and to known human pathogens. Finally, we evaluated the effects of ectoparasite load on the likelihood of infection with bartonellae and hemoplasmas in these bats.

## Methods

### Capture and sampling of bats

We sampled *M. condylurus*, *M. pumilus*, and *C. cor* during May through June of 2023 and June of 2024 in Taita-Taveta County, Kenya. Taita-Taveta, similar to regions across Sub-Saharan Africa, is comprised of fragmented highlands and lower-elevation grasslands and woodlands with rural villages that rely on subsistence farming (Abera *et al*. 2022). While Taita-Taveta county is highly biodiverse, urbanization over the last decades has resulted in dramatic forest and grassland loss (Nyongesa *et al*. 2022).

Bats were trapped with mist nets while exiting building roosts or flying over a body of water across 21 sites. Bats were placed in individual cotton bags and taken to a secure area at a nearby research station for processing. During the sampling process, we collected approximately 10 µL of blood from the cephalic vein of each bat for storage on Whatman FTA cards (QIAGEN, Hilden, Germany) for bacterial pathogen screening. We also visually inspected each bat for ectoparasites (mites, fleas, ticks, and bat flies) and recorded overall ectoparasite load (as an index: 0, 1–5, 6–10, 11–15, 16+). We collected representative ectoparasites from each bat (when available) using tweezers and stored them in 96% ethanol. Bats were then released at night near the site of capture following processing. Field procedures were performed according to guidelines for the safe and humane handling of bats published by of the American Society of Mammalogists (Sikes *et al*. 2016) and were approved by the Institutional Animal Care and Use Committees of the University of Arkansas (2403002206). Fieldwork and sampling were authorized by the Kenyan Wildlife Service under permits (WRTI-0100-10-21, KWS/BRM/5001).

### Ectoparasite identification

Prior to identification, mites were cleared in 5% lactic acid, mounted on glass slides using Hoyer’s medium, and incubated for one week at 50°C to ensure proper setting. We identified the bat mites and bat fleas under a compound microscope (100-400x magnification) based on morphological characteristics (mites: number and placement of setae on dorsal shield, prodonotal region, and opisthonotal region; fleas: genal teeth, number and placement of setae, length of pronotum) using taxonomic keys, narrowed down by previous records of ectoparasites on *M. condylurus* and *M. pumilus* (Smit 1957; Till & Evans 1966). We identified argasid ticks and nycteribiid bat flies with environmental scanning electron microscopy images based on morphological characteristics (ticks: body size, integument, margins, mouthparts; bat flies: microtrichia on sternal plate, theca width, posterior margin cavities, gular setae) using taxonomic keys, cross checked with previous records of ectoparasites described on *C. cor* (Hoogstraal 1955; Shapiro *et al*. 2016).

### DNA extraction, PCR amplification, and amplicon sequencing

Genomic DNA was extracted from bat blood on Whatman FTA cards using Zymo *Quick*-DNA Tissue/Insect Microprep Kits (Zymo Research, Irvine, CA, USA), following the manufacturer’s instructions. Ectoparasites collected from each bat were sorted by taxa (mites, fleas, ticks, and bat flies). All ectoparasites were pooled by ectoparasite taxon and by individual bat, and crushed. DNA from extracted using a non-commercial 5% (m/v)100 mesh Chelex® resin protocol (Bio-Rad Laboratories, Inc, Hercules, CA, USA) following a modified version of McMicheal *et al*. (2011). Briefly, Proteinase K (20mg/mL) was added to each sample, followed by an overnight incubation at 55°C prior to being heated at 90°C for 10 minutes. Following centrifugation, approximately 100 μL of the resulting supernatant (containing DNA) was recovered.

We performed nested PCR and gel electrophoresis to determine the presence of *Bartonella* spp. (targeting the partial citrate synthase [*gltA*] gene) and conventional PCR and gel electrophoresis to determine the presence of hemoplasmas (targeting the partial 16S ribosomal RNA gene) in both bat blood and ectoparasite DNA. Primers and procedures used for both PCR assays have been described and validated previously (Volokhov *et al*. 2011; Becker *et al*. 2025). We included extraction controls (blank FTA card punches), negative controls (nuclease-free water), and positive controls from prior samples in all PCRs. *Bartonella* spp. amplicons (∼300 bp) and hemoplasma amplicons (∼850–900 bp) were identified during gel electrophoresis. A subset of the positive amplicons were extracted from gels and purified using Zymoclean Gel DNA Recovery Kits (Zymo Research, Irvine, CA, USA). Sanger sequencing of purified amplicons was performed by Eurofins Genomics (Louisville, KY, USA).

### Statistical analysis

To evaluate the influence of bat ectoparasite load, sex, age class, and year on the odds of infection with hemoplasmas or bartonellae, we used generalized linear mixed models (GLMM) fit separately for *M. condylurus* and *M. pumilus,* with site included as a random intercept in each model. We used generalized linear models (GLM) to evaluate the relationship between these four predictors on the likelihood of both infections for *C. cor,* since this bat species was sparsely distributed across sites. All models were fitted with a binomial error distribution (logit link) using the glmer() function in the *lme4* package (Bates *et al*. 2015) and glm() function. To assess model performance, we utilized the *performance* package (Lüdecke *et al*. 2021) to calculate marginal and conditional coefficients of determination (R^2^) for GLMMs and a coefficient of discrimination (Pseudo R^2^) for GLMs. Statistical analyses were performed in R (version 4.4.3; R Core Team 2026).

### Phylogenetic analysis

Chromatograms for forward and reverse strands for partial *gltA* and partial 16S rRNA sequences were visually inspected and edited for quality using Chromas v.2.6.6 (Technelysium Pty Ltd). Consensus sequences were made by assembling the forward and reverse sequences in MEGA11 (Tamura *et al*. 2021). NCBI BLASTn was then used to identify related *Bartonella gltA* (*Bartonella)* and 16S rRNA (hemoplasma) sequences, which were aligned with the sequences from this study using MUSCLE. Phylogenetic reconstruction was performed using IQ-TREE (Minh *et al*. 2020) using a GTR + I + G model. To assess node support, we performed 1,000 replicates of Ultrafast Bootstrap approximation.

## Results

We sampled a total of 767 bats across 21 sites, including 374 *M. condylurus*, 320 *M. pumilus*, and 73 *C. cor*. Of the 21 sites, *M. condylurus* was present at 10 sites (47.6%), *M. pumilus* was present at 17 sites (81.0%), and *C. cor* was present at five sites (23.8%). *M. condylurus* and *M. pumilus* were both present at nine of the 21 sites (43.0%), and *M. pumilus* and *C. cor* were both present at two of the 21 sites (9.5%).

We identified one species of flea, *Lagaropsylla idae*, and one species of mite*, Chelanyssus aethiopicus,* on both *M. condylurus* and *M. pumilus* (Fig. S1). Additionally, we found one species of tick, *Argas boueti*, and one species of bat fly, *Raymondia planiceps*, on *C. cor* (Fig. S1). All ectoparasites were previously described on the host species from this study (Hoogstraal 1955; Smit 1957; Till & Evans 1966; Shapiro *et al*. 2016). Ectoparasites generally were observed on 78.5% of bats. Specifically, 33.4% (N_pooled_=125) of *M. condylurus* had at least one flea and 92.0% (N_pooled_=344) of *M. condylurus* had at least one mite (Table 1). For *M. pumilus,* 6.9% (N_pooled_=22) had at least one flea and 61.6% (N_pooled_=197) had at least one mite (Table 1). For *C. cor,* 50.7% (N_pooled_=37) had at least one tick and 5.5% (N_pooled_=4) had at least one bat fly (Table 1).

**Table 1.** Individual-host pooled prevalence of *Bartonella* infection across ectoparasites species collected from synanthropic bats in Taita-Taveta County, Kenya. No ectoparasite pools were positive for hemoplasmas.

| Species | Host | N<sub>pooled</sub><br>groups | Individual-host pooled<br>*Bartonella* spp.<br>prevalence (95% CI) |
| --- | --- | --- | --- |
| **Flea** (*Lagaropsylla*<br>*idea*) | *Mops condylurus* | 125 | 24.0% (16.8–32.5%) |
|  | *Mops pumilus* | 22 | 27.3% (10.7–50.2%) |
|  | *M. condylurus* & *M. pumilus* | 147 | 24.5% (17.8–32.3%) |
| **Mite** (*Chelanyssus*<br>*aethiopicus*) | *Mops condylurus* | 344 | 21.3% (15.8–27.7%) |
|  | *Mops pumilus* | 197 | 19.8% (15.7–24.3%) |
|  | *M. condylurus* & *M. pumilus* | 541 | 20.3% (17.0–23.9%) |
| **Tick** (*Argas boueti*) | *Cardioderma cor* | 37 | 13.5% (4.5–28.8%) |
| **Bat fly** (*Raymondia*<br>*planiceps*) | *Cardioderma cor* | 4 | 25.0% (0.6–80.6%) |

Overall, 17.9% of all bats across all sites were *Bartonella* spp. positive, with 24.6% prevalence in *M. condylurus*, 5.3% prevalence in *M. pumilus*, and 38.4% in *C. cor* (Table 2). Overall, 19.3% of all bats across all sites were hemoplasma positive, with 18.7% prevalence in *M. condylurus*, 20.0% prevalence in *M. pumilus*, and 19.2% in *C. cor* (Table 2). For ectoparasites, 24.5% of flea pools (pooled by bat; N_pooled_=147), 20.3% of mite pools (pooled by bat; N_pooled_=541), 13.5% of tick pools (pooled by bat; N_pooled_=37), and 25.0% of bat fly pools (pooled by bat; N_pooled_=4) were positive for bartonellae (Table 1). We did not detect hemoplasmas in any of the ectoparasites.

**Table 2.** Bacterial pathogen prevalence in the three most common synanthropic bats in Taita-Taveta County, Kenya.

| Species | *Bartonella* spp. prevalence<br>(95% CI) | Hemoplasma prevalence<br>(95% CI) |
| --- | --- | --- |
| *Mops condylurus* (N=375) | 24.6% (20.3–29.3%) | 18.7% (14.9–23.0%) |
| *Mops pumilus* (N=321) | 5.3% (3.1–8.4%) | 20.0% (15.8–24.8%) |
| *Cardioderma cor* (N=74) | 38.4% (27.2–50.4%) | 19.2% (10.9–30.1%) |
| Total (N=767) | 17.9% (15.3–20.8%) | 19.3% (16.6–22.3%) |

Ectoparasite load, sex, age class, and year had no effect on *Bartonella* spp. infection in *M. condylurus* or *M. pumilus* (Table S1). For these GLMMs, site explained 0.0% and 10.7% of the variance in infection risk in *M. condylurus* and *M. pumilus,* respectively. However, *Bartonella* spp. infection in *C. cor* was positively associated with higher ectoparasite load (OR=2.93; 95% CI:1.60–6.23; p<0.01; Fig. 1) and negatively associated with being male (OR=0.15; 95% CI:0.02–0.79; p<0.05). Age class and year had no effect on *Bartonella* spp. infection in *C. cor* (age: OR<0.01, 95% CI: 0–Inf; p=0.99; year: OR=1.52, 95% CI: 0.31–7.82; p=0.60), and these predictors (ectoparasite load, sex, age, year) overall explained 33.7% of the variance in infection risk. Likewise, ectoparasite load, sex, age class, and year had no effect on hemoplasma infection for each bat species (Table S1). For these GLMMs, site explained 0.4% of the variance in infection risk for *M. condylurus* and 16.7% for *M. pumilus*.

**Figure 1.**
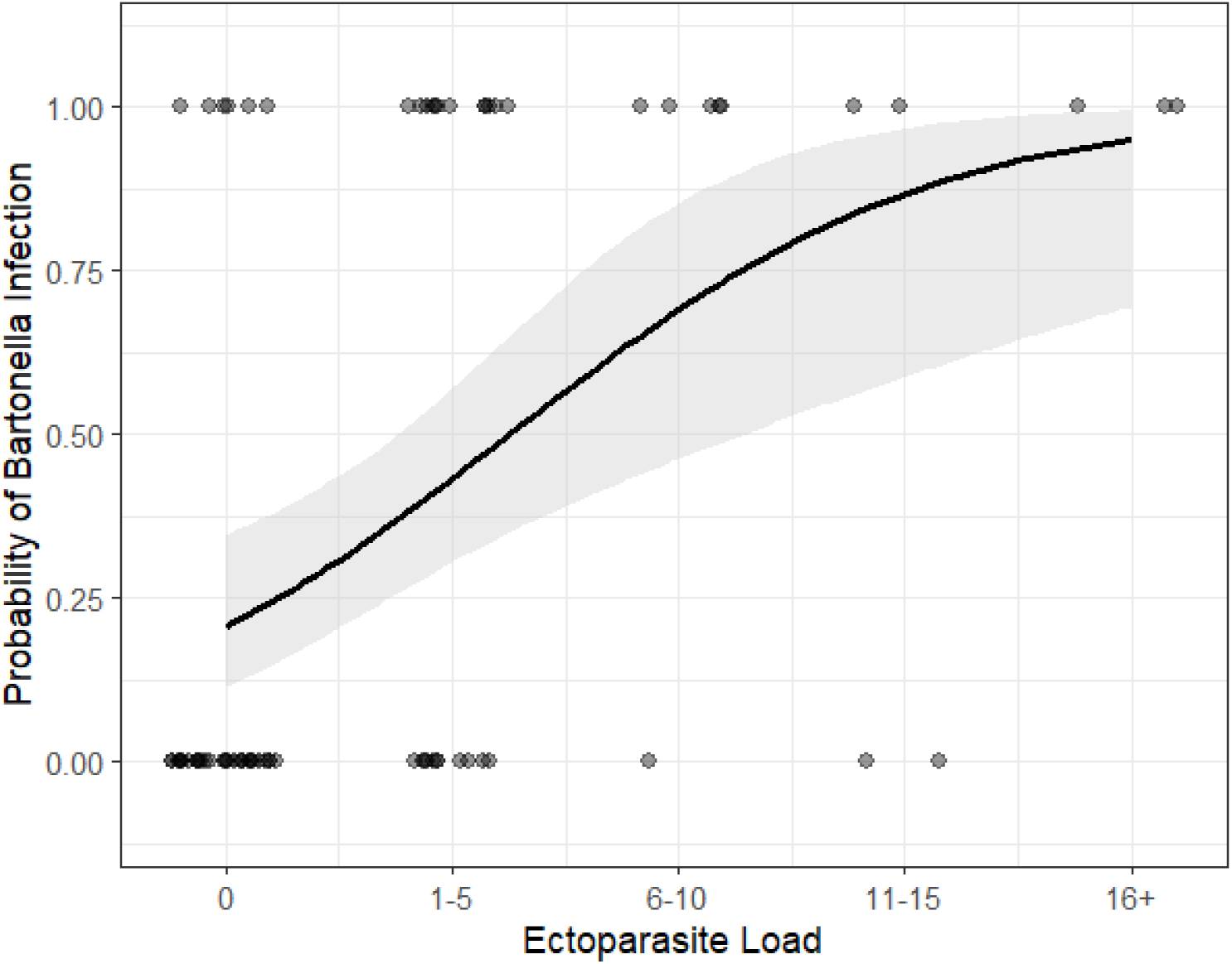
Predicted probability of *Bartonella* spp. infection in *Cardioderma cor* as a function of ectoparasite load. Ectoparasite load was treated as a continuous predictor using numeric ranks corresponding to binned ectoparasite load. The solid line represents the predicted logistic regression curve with shaded 95% confidence interval, overlaid on jittered raw infection data.

Phylogenetic analysis of the *gltA* gene revealed a high diversity of *Bartonella* lineages circulating within bats and their ectoparasites (Fig. 2). Unlike the hemoplasma phylogeny, which showed distinct lineages corresponding to host taxonomy, *Bartonella* sequences demonstrated phylogenetic overlap between different bat species and between hosts and their vectors. Within this phylogeny, we identified multiple supported clades where sequences from bats and their ectoparasites are highly conserved, as seen in *M. condylurus* and their fleas and *C. cor* and their ticks and bat flies (99.10%–100.00% identical; >90% bootstrap support; Fig. 2).

**Figure 2.**
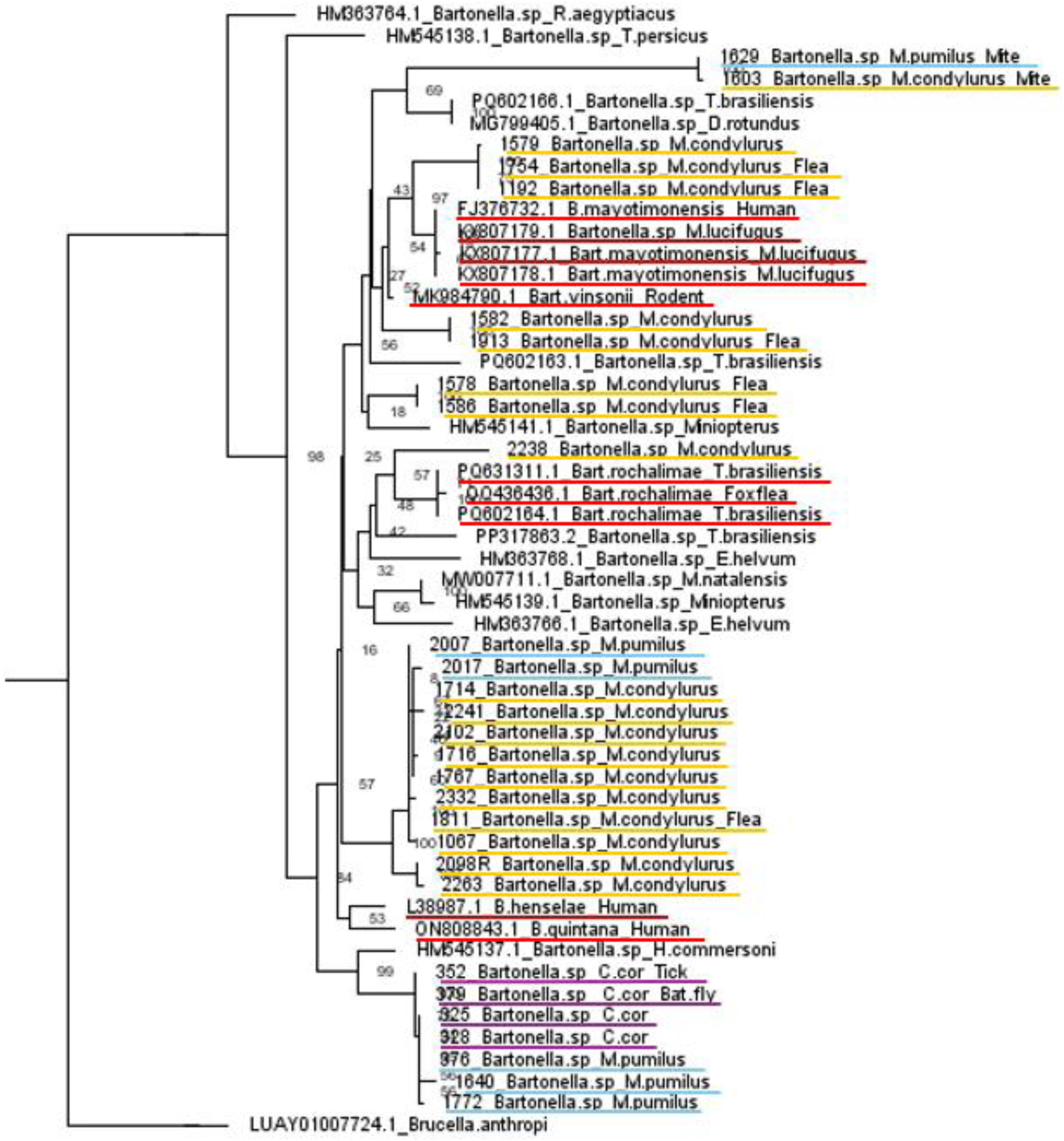
Phylogenetic relationships among sequences of *Bartonella* spp. partial citrate synthase gene (*gltA*) detected in *M. condylurus* (yellow), *M. pumilus* (blue), and *C. cor* (purple), and their ectoparasites from this study, as well as those previously described. Known zoonotic pathogens are underlined in red. Nodes display bootstrap support based on 1000 randomizations.

Phylogenetic analysis of the partial 16S rRNA gene revealed that hemoplasma sequences clustered into distinct lineages that correspond to host taxonomy (Fig. 3). The hemoplasma sequences from *M. condylurus* and *M. pumilus* were highly conserved, with 98.82%–99.62% sequence identity. These sequences formed a supported, monophyletic clade (88% bootstrap support; Fig. 2), indicating a shared bacterial lineage within these molossid bats (*M. condylurus* and *M. pumilus*). The hemoplasma sequences identified in *M. condylurus* and *M. pumilus* also grouped closely with lineages previously described in New World molossids, including *Tadarida brasiliensis* and *Molossus nigricans* (80% bootstrap support; Fig. 3). The sequences from *C. cor* were phylogenetically distant from the molossid-associated clade and instead clustered within a lineage containing human-associated hemoplasmas, previously identified in *Miniopterus schreibersii* in Spain with 97.85% sequence identity (91% bootstrap support; Fig. 3).

**Figure 3.**
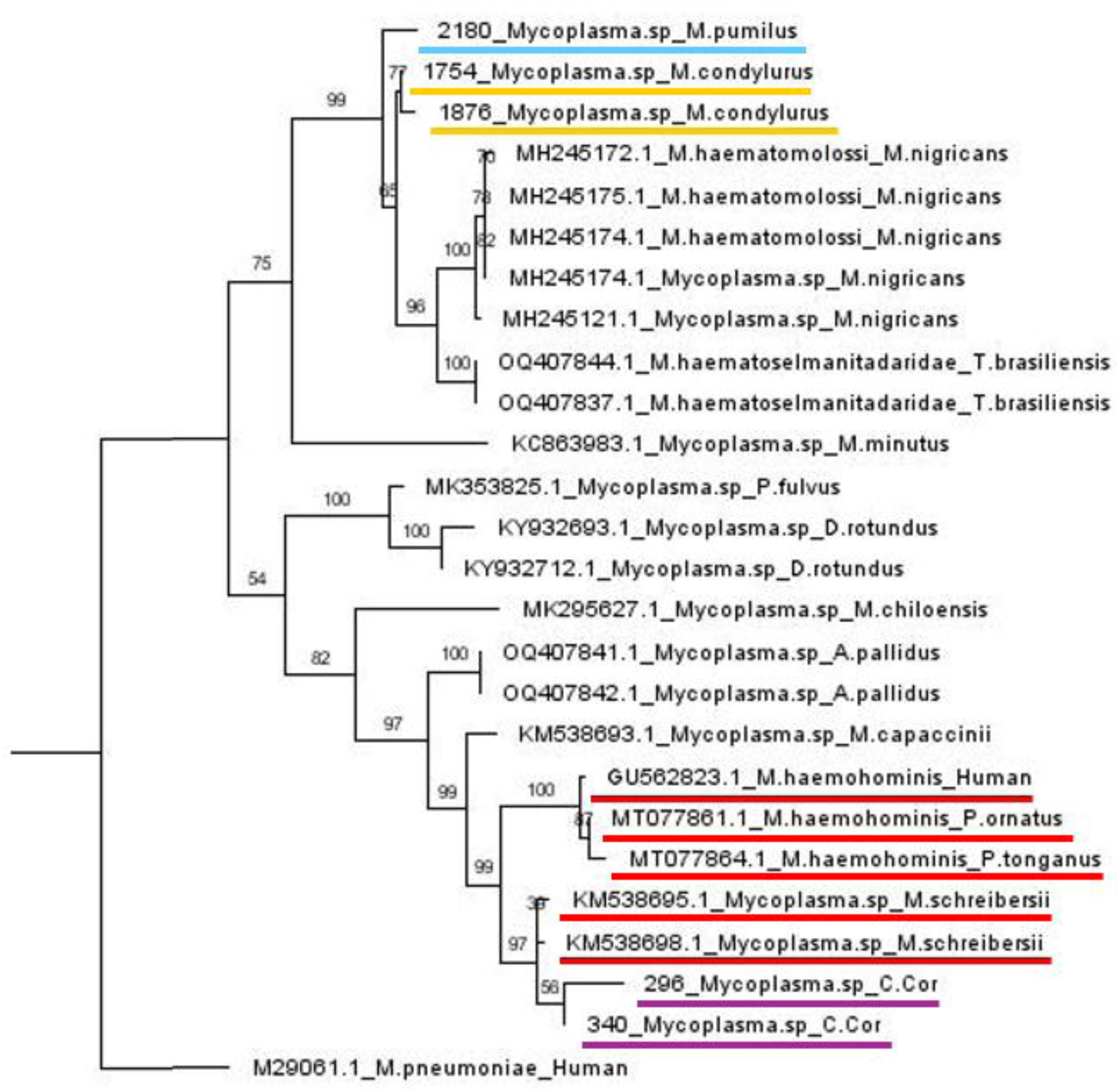
Phylogenetic relationships among sequences of hemoplasma partial 16S rRNA gene detected in *M. condylurus* (yellow), *M. pumilus* (blue), and *C. cor* (purple), as well as those from previously described *Mycoplasma* spp. Known zoonotic pathogens are underlined in red. Nodes display bootstrap support based on 1000 randomizations.

## Discussion

Our results demonstrate that synanthropic Kenyan bats host diverse bacterial pathogens, with bartonellae demonstrating broad transmission across multiple species and vectors and hemoplasmas instead demonstrating host specialization. Importantly, we found evidence that *C. cor* hosts a hemoplasma that is closely related to human-associated lineages. Overall, we show that synanthropic bats host bacteria of potential public health concern, highlighting the need to investigate the emerging impacts of these pathogens on human health in Kenya and elsewhere in Sub-Saharan Africa.

*Bartonella* sequences showed minimal host specialization, with a diverse range of hosts and vectors sharing similar genotypes and forming multi-host clades. This lack of host specificity suggests *Bartonella* lineages are shared between species, such as between *M. pumilus* and *C. cor,* potentially mediated by a shared environment. Previous studies assessing the relationship between evolutionary and ecological similarities of host species and bacterial pathogens have also found that closely related *Bartonella* genotypes are shared across taxonomically diverse species (Lei & Olival 2014; McKee *et al*. 2019; Becker *et al*. 2024). In contrast, the clustering of hemoplasma sequences from the two species of molossid bats in our study and previously described New World molossids suggest strong host specialization within the family Molossidae. Our findings align with previous research that found hemoplasmas show strong host specificity (Becker *et al*. 2020; Millán *et al*. 2021). Our investigation of bat-borne hemoplasma expands the known geographic and host range of hemoplasmas in Africa, which is largely understudied compared to other regions (Di Cataldo *et al*. 2020; Millán *et al*. 2021).

The detection of a hemoplasma in *C. cor* that is closely related to a lineage found in *Miniopterus schreibersii* is particularly significant, given previous detection of closely related hemoplasmas in immunocompromised children in Spain (Esperon *et al*. 2025). The closely related hemoplasmas in the East African *C. cor* and the European *M. schreibersii* indicate a broader geographical distribution and greater host plasticity than previously documented (Millan *et al*. 2015; Esperon *et al*. 2025), as *C. cor* and *M. schreibersii* do not have overlapping ranges and are in distinct taxonomic suborders (Csada 1996; Kohl & Kurth 2014). Similar to these results, a previous African survey on bat-borne hemoplasmas found evidence for cross-species transmission between taxonomically distant bats (Di Cataldo *et al*. 2020). While these findings highlight a potential zoonotic risk, our current characterization of the *C. cor* hemoplasma genotype is limited to the partial 16S rRNA gene. Sequencing additional genes, or establishing whole genomes, will be required to confirm the relatedness between the *C. cor* lineage and zoonotic hemoplasma lineages.

The lack of association between ectoparasite load and infection status in most of these bat species, in combination with low prevalences in ectoparasites, suggests that vector–mediated transmission may not be the primary driver of pathogen maintenance in these populations, particularly for hemoplasmas. Instead, a combination of multiple transmission routes may be responsible (Cohen et al. 2018). While bat flies have been linked to *Bartonella* transmission in both captive and wild populations of bats (Fagre *et al*. 2023; McKee *et al*. 2026), alternative transmission pathways, such as direct transmission via aggressive interactions, have been documented from other animals (Keret *et al*. 1998; Windsor *et al*. 2001). For hemoplasmas, transmission routes remain largely unknown, although suggested routes include vector-borne, blood-borne, or vertical transmission (Tasker 2020). The lack of hemoplasma positive ectoparasites in this study, despite high hemoplasma prevalence in bats, supports direct transmission between bats as a primary route of hemoplasma transmission in these populations, which is consistent with observations from a rodent-borne hemoplasma study (Cohen *et al*. 2018).

We found some evidence that bat hosts and their ectoparasites share closely related *Bartonella* genotypes, indicating that arthropod vectors may be at least partially responsible for transmission of bacteria within roosts (Brook *et al*. 2015; Arendt *et al*. 2024). However, confirming whether ectoparasites are functional vectors is diagnostically challenging, due to the difficulty of distinguishing between biological infection and mechanical carriage of remnant host blood (Kapustová *et al*. 2025). Furthermore, the lack of hemoplasma-positive ectoparasites in this study, could be partly attributed to digestive processes in ectoparasites that quickly break down blood meals and associated pathogens to varying degrees, making it difficult to detect pathogens in some cases, even when present (Zellner *et al*. 2026).

Our study identified high infection prevalence and pathogen sharing of bacterial pathogens within anthropogenic habitats. We found that bacterial pathogens followed distinct phylogenetic patterns occurring in the same synanthropic environments, with multi-host bartonellae lineages and host-specific molossid hemoplasma lineages. Additionally, we detected a potentially zoonotic pathogen in a synanthropic bat species. This study represents only the second investigation of bat-borne hemoplasmas in Africa. Consequently, expanded surveillance is essential to determine the geographic distribution, phylogenetic diversity, and zoonotic implications of both bartonellae and hemoplasmas in Africa. Understanding bacterial pathogen dynamics at the human–wildlife interface is critical to developing public health strategies that protect people from potential outbreaks.

## Acknowledgments

We would like to thank the Taita people for their time and partnership with this project. We also thank the Taita Environmental Research and Resource Arc for their facilities and assistance. We acknowledge and thank Hayden McGee, Anna Bolding, Raven Leggett, and Annabelle Erisman for assistance with laboratory assays.

## References

Abera, T. A., Vuorinne, I., Munyao, M., Pellikka, P. K. E., & Heiskanen, J. (2022). Land Cover Map for Multifunctional Landscapes of Taita Taveta County, Kenya, Based on Sentinel-1 Radar, Sentinel-2 Optical, and Topoclimatic Data. Data, 7(3), 36. 10.3390/data7030036

Alcorn, K., Gerrard, J., Cochrane, T., Graham, R., Jennison, A., Irwin, P. J., & Barbosa, A. D. (2021). First Report of Candidatus Mycoplasma haemohominis Infection in Australia Causing Persistent Fever in an Animal Carer. Clinical Infectious Diseases: An Official Publication of the Infectious Diseases Society of America, 72(4), 634–640. 10.1093/cid/ciaa089

Allocati, N., Petrucci, A. G., Di Giovanni, P., Masulli, M., Di Ilio, C., & De Laurenzi, V. (2016). Bat–man disease transmission: Zoonotic pathogens from wildlife reservoirs to human populations. Cell Death Discovery, 2(1), 16048. 10.1038/cddiscovery.2016.48

Arendt, M., Stadler, J., Ritzmann, M., Ade, J., Hoelzle, K., & Hoelzle, L. E. (2024). Hemotrophic Mycoplasmas—Vector Transmission in Livestock. Microorganisms, 12(7), 1278. 10.3390/microorganisms12071278

Bai, Y., Osinubi, M. O. V., Osikowicz, L., McKee, C., Vora, N. M., Rizzo, M. R., Recuenco, S., Davis, L., Niezgoda, M., Ehimiyein, A. M., Kia, G. S. N., Oyemakinde, A., Adeniyi, O. S., Gbadegesin, Y. H., Saliman, O. A., Ogunniyi, A., Ogunkoya, A. B., & Kosoy, M. Y. (2018). Human Exposure to Novel Bartonella Species from Contact with Fruit Bats—Volume 24, Number 12—December 2018—Emerging Infectious Diseases journal—CDC. 10.3201/eid2412.181204

Bates, D., Mächler, M., Bolker, B., Walker, S. (2015). Fitting Linear Mixed-Effects Models Using lme4. Journal of Statistical Software, 67(1), 1–48. doi:10.18637/jss.v067.i01.

Becker, D. J., Bergner, L. M., Bentz, A. B., Orton, R. J., Altizer, S., & Streicker, D. G. (2018). Genetic diversity, infection prevalence, and possible transmission routes of Bartonella spp. In vampire bats. PLOS Neglected Tropical Diseases, 12(9), e0006786. 10.1371/journal.pntd.0006786

Becker, D. J., Speer, K. A., Brown, A. M., Fenton, M. B., Washburne, A. D., Altizer, S., Streicker, D. G., Plowright, R. K., Chizhikov, V. E., Simmons, N. B., & Volokhov, D. V. (2020). Ecological and evolutionary drivers of haemoplasma infection and bacterial genotype sharing in a Neotropical bat community. Molecular Ecology, 29(8), 1534–1549. 10.1111/mec.15422

Becker, D. J., Dyer, K. E., Lock, L. R., Olbrys, B. L., Pladas, S. A., Sukhadia, A. A., Demory, B., Nunes Batista, J. M., Pineda, M., Simmons, N. B., Adams, A. M., Frick, W. F., O’Mara, M. T., & Volokhov, D. V. (2024). Geographically widespread and novel hemotropic mycoplasmas and *Bartonella*e in Mexican free-tailed bats and sympatric North American bat species. mSphere, 10(1), e00116–24. 10.1128/msphere.00116-24

Betke B.A., Gottdenker N.L., Meyers L.A., Becker D.J. (2024). Ecological and evolutionary characteristics of anthropogenic roosting ability in bats of the world. Iscience, 19;27(7).

Brandell, E. E., Becker, D. J., Sampson, L., & Forbes, K. M. (2021). Demography, education, and research trends in the interdisciplinary field of disease ecology. Ecology and Evolution, 11(24), 17581–17592. 10.1002/ece3.8466

Brook, C. E., Bai, Y., Dobson, A. P., Osikowicz, L. M., Ranaivoson, H. C., Zhu, Q., Kosoy, M. Y., & Dittmar, K. (2015). Bartonella spp. In Fruit Bats and Blood-Feeding Ectoparasites in Madagascar. PLOS Neglected Tropical Diseases, 9(2), e0003532. 10.1371/journal.pntd.0003532

Buchmann, C. M., Schurr, F. M., Nathan, R., & Jeltsch, F. (2013). Habitat loss and fragmentation affecting mammal and bird communities—The role of interspecific competition and individual space use. Ecological Informatics, The Analysis and Application of Spatial Ecological Data to Support the Conservation of Biodiversity, 14, 90–98. 10.1016/j.ecoinf.2012.11.015

Chomel, B. B., Stuckey, M. J., Boulouis, H.-J., & Aguilar-Setién, A. (2014). Bat-Related Zoonoses. Zoonoses - Infections Affecting Humans and Animals, 697–714. 10.1007/978-94-017-9457-2_28

Cohen C., Shemesh M., Garrido M., Messika I., Einav M., Khokhlova I., Tasker S., Hawlena H. Haemoplasmas in wild rodents: Routes of transmission and infection dynamics. Molecular Ecology. 2018 Sep;27(18):3714–26.

Csada, R. (1996). Cardioderma cor. Mammalian Species, 519: 1–4.

Descloux, E., Mediannikov, O., Gourinat, A.-C., Colot, J., Chauvet, M., Mermoud, I., Desoutter, D., Cazorla, C., Klement-Frutos, E., Antonini, L., Levasseur, A., Bossi, V., Davoust, B., Merlet, A., Goujart, M.-A., Oedin, M., Brescia, F., Laumond, S., Fournier, P.-E., & Raoult, D. (2021). Flying Fox Hemolytic Fever, Description of a New Zoonosis Caused by Candidatus Mycoplasma haemohominis. Clinical Infectious Diseases: An Official Publication of the Infectious Diseases Society of America, 73(7), e1445–e1453. 10.1093/cid/ciaa1648

Di Cataldo, S., Kamani, J., Cevidanes, A., Msheliza, E. G., & Millán, J. (2020). Hemotropic mycoplasmas in bats captured near human settlements in Nigeria. Comparative Immunology, Microbiology and Infectious Diseases, 70, 101448. 10.1016/j.cimid.2020.101448

Dietrich, M., Tjale, M. A., Weyer, J., Kearney, T., Seamark, E. C. J., Nel, L. H., Monadjem, A., & Markotter, W. (2016). Diversity of Bartonella and Rickettsia spp. In Bats and Their Blood-Feeding Ectoparasites from South Africa and Swaziland. PLOS ONE, 11(3), e0152077. 10.1371/journal.pone.0152077

Ecke, F., Han, B. A., Hörnfeldt, B., Khalil, H., Magnusson, M., Singh, N. J., & Ostfeld, R. S. (2022). Population fluctuations and synanthropy explain transmission risk in rodent-borne zoonoses. Nature Communications, 13(1), 7532. 10.1038/s41467-022-35273-7

Entwistle A, Racey P, Speakman J. Roost Selection by the Brown Long-Eared Bat Plecotus auritus. The Journal of Applied Ecology. 1997 Apr 1;34:399. doi:10.2307/2404885

Eremeeva M.E., Gerns H.L., Lydy S.L., Goo J.S., Ryan E.T., Mathew S.S., Ferraro M.J., Holden J.M., Nicholson W.L., Dasch G.A., Koehler J.E. (2007). Bacteremia, fever, and splenomegaly caused by a newly recognized Bartonella species. New England Journal of Medicine, 7;356(23):2381–7.

Esperón, F., Martín-Maldonado, B., Iglesias, I., Neves, E., Seri, E., García-Sanchez, P., Morillas-Mingorance, Á., & Méndez-Echevarría, A. (2025). Bat-Associated Hemotropic Mycoplasmas in Immunosuppressed Children, Spain, 2024. Emerging Infectious Diseases, 31(10), 1980–1983. 10.3201/eid3110.250862

Fagre, A. C., Islam, A., Reeves, W. K., Kading, R. C., Plowright, R. K., Gurley, E. S., & McKee, C. D. (2023). Bartonella Infection in Fruit Bats and Bat Flies, Bangladesh. Microbial Ecology, 86(4), 2910–2922. 10.1007/s00248-023-02293-9

Friant, S., Mistrick, J., Luis, A. D., Harden, C., Simons, D., Fichet-Calvet, E., Gibb, R., Grube, N., Henttonen, H., Imirizian, N., Moses, L., Perry, G. H., Redding, D., Stenseth, N. C., Vandegrift, K., Bjornstad, O. N., Dobson, A., Lloyd-Smith, J. O., & Hudson, P. J. (2025). Reducing the threats of rodent-borne zoonoses requires an understanding and leveraging of three key pillars: Disease ecology, synanthropy, and rodentation. The Lancet Planetary Health, 9(9). 10.1016/j.lanplh.2025.101300

Gravinatti, M. L., Barbosa, C. M., Soares, R. M., & Gregori, F. (2020). Synanthropic rodents as virus reservoirs and transmitters. Revista Da Sociedade Brasileira de Medicina Tropical, 53, e20190486. 10.1590/0037-8682-0486-2019

Hassell, J. M., Begon, M., Ward, M. J., & Fèvre, E. M. (2017). Urbanization and Disease Emergence: Dynamics at the Wildlife-Livestock-Human Interface. Trends in Ecology & Evolution, 32(1), 55–67. 10.1016/j.tree.2016.09.012

Hattori, N., Kuroda, M., Katano, H., Takuma, T., Ito, T., Arai, N., Yanai, R., Sekizuka, T., Ishii, S., Miura, Y., Tokunaga, T., Watanabe, H., Nomura, N., Eguchi, J., Hasegawa, H., Nakamaki, T., Wakita, T., & Niki, Y. (2020). Candidatus Mycoplasma haemohominis in Human, Japan. Emerging Infectious Diseases, 26(1), 11–19. 10.3201/eid2601.190983

Jackson, R. T., Webala, P. W., Ogola, J. G., Lunn, T. J., & Forbes, K. M. (2023). Roost selection by synanthropic bats in rural Kenya: Implications for human–wildlife conflict and zoonotic pathogen spillover. Royal Society Open Science, 10(9), 230578. 10.1098/rsos.230578

Kamani, J., Baneth, G., Mitchell, M., Mumcuoglu, K. Y., Gutiérrez, R., & Harrus, S. (2014). Bartonella Species in Bats (Chiroptera) and Bat Flies (Nycteribiidae) from Nigeria, West Africa. Vector Borne and Zoonotic Diseases, 14(9), 625–632. 10.1089/vbz.2013.1541

Kapustová, A., Kulich Fialová, M., Svobodová, M., & Brzoňová, J. (2025). Combining blood meal analysis and parasite detection yields a more comprehensive understanding of insect host feeding patterns. Parasites & Vectors, 18(1), 288. 10.1186/s13071-025-06931-8

Keret, D., Giladi, M., Kletter, Y., & Wientroub, S. (1998). Cat-scratch disease osteomyelitis from a dog scratch. The Journal of Bone and Joint Surgery. British Volume, 80(5), 766–767.

Kohl, C., & Kurth, A. (2014). European Bats as Carriers of Viruses with Zoonotic Potential. Viruses, 63390, 3110–3128. 10.3390/v6083110

Kosoy, M., Bai, Y., Lynch, T., Kuzmin, I. V., Niezgoda, M., Franka, R., Agwanda, B., Breiman, R. F., & Rupprecht, C. E. (2010). *Bartonella* spp. In bats, Kenya. Emerging Infectious Diseases, 16(12), 1875–1881. 10.3201/eid1612.100601

Lagaropsylla idae. (n.d.). Retrieved April 4, 2026, from https://insecta.pro/taxonomy/797785

Lei, B. R., & Olival, K. J. (2014). Contrasting Patterns in Mammal–Bacteria Coevolution: *Bartonella* and Leptospira in Bats and Rodents. PLoS Neglected Tropical Diseases, 8(3), e2738. 10.1371/journal.pntd.0002738

Letko, M., Seifert, S. N., Olival, K. J., Plowright, R. K., & Munster, V. J. (2020). Bat-borne virus diversity, spillover and emergence. Nature Reviews Microbiology, 18(8), 461–471. 10.1038/s41579-020-0394-z

Lin, E. Y., Tsigrelis, C., Baddour, L. M., Lepidi, H., Rolain, J.-M., Patel, R., & Raoult, D. (n.d.). Candidatus Bartonella mayotimonensis and Endocarditis—Volume 16, Number 3—March 2010—Emerging Infectious Diseases journal—CDC. 10.3201/eid1603.081673

Lüdecke, D., Ben-Shachar, M., Patil, I., Waggoner, P., Makowski, D. (2021). performance: An R Package for Assessment, Comparison and Testing of Statistical Models. Journal of Open Source Software, 6(60), 3139. doi:10.21105/joss.03139.

Markotter, W., Coertse, J., De Vries, L., Geldenhuys, M., & Mortlock, M. (2020). Bat-borne viruses in Africa: A critical review. Journal of Zoology (London, England : 1987), 311(2), 77–98. 10.1111/jzo.12769

McFarlane, R., Sleigh, A., & McMichael, T. (2012). Synanthropy of Wild Mammals as a Determinant of Emerging Infectious Diseases in the Asian–Australasian Region. Ecohealth, 9(1), 24–35. 10.1007/s10393-012-0763-9

McKee, C. D., Krawczyk, A. I., Sándor, A. D., Görföl, T., Földvári, M., Földvári, G., Dekeukeleire, D., Haarsma, A.-J., Kosoy, M. Y., Webb, C. T., & Sprong, H. (2019). Host Phylogeny, Geographic Overlap, and Roost Sharing Shape Parasite Communities in European Bats. Frontiers in Ecology and Evolution, 7. 10.3389/fevo.2019.00069

McKee C.D., Webb C.T., Kosoy M.Y., Suu-Ire R, Ntiamoa-Baidu Y., Cunningham A.A., Wood J.L., Hayman D.T. (2026). Manipulating vector transmission reveals local processes in Bartonella communities of bats. Parasitology, 2:1–40.

McMichael, G. L., Highet, A. R., Gibson, C. S., Goldwater, P. N., O’Callaghan, M. E., Alvino, E. R., & MacLennan, A. H. (2011). Comparison of DNA Extraction Methods from Small Samples of Newborn Screening Cards Suitable for Retrospective Perinatal Viral Research. Journal of Biomolecular Techniques : JBT, 22(1), 5–9.

MEGA11: Molecular Evolutionary Genetics Analysis Version 11 | Molecular Biology and Evolution | Oxford Academic. (n.d.). Retrieved April 5, 2026, from https://academic.oup.com/mbe/article/38/7/3022/6248099

Messick, J. B. (2004). Hemotrophic mycoplasmas (hemoplasmas): A review and new insights into pathogenic potential. Veterinary Clinical Pathology, 33(1), 2–13. 10.1111/j.1939-165x.2004.tb00342.x

Millán, J., Di Cataldo, S., Volokhov, D. V., & Becker, D. J. (2021). Worldwide occurrence of haemoplasmas in wildlife: Insights into the patterns of infection, transmission, pathology and zoonotic potential. Transboundary and Emerging Diseases, 68(6), 3236–3256. 10.1111/tbed.13932

Millán, J., López-Roig, M., Delicado, V., Serra-Cobo, J., & Esperón, F. (2015). Widespread infection with hemotropic mycoplasmas in bats in Spain, including a hemoplasma closely related to “Candidatus Mycoplasma hemohominis.” Comparative Immunology, Microbiology and Infectious Diseases, 39, 9–12. 10.1016/j.cimid.2015.01.002

Minh, B. Q., Schmidt, H. A., Chernomor, O., Schrempf, D., Woodhams, M. D., von Haeseler, A., & Lanfear, R. (2020). IQ-TREE 2: New Models and Efficient Methods for Phylogenetic Inference in the Genomic Era. Molecular Biology and Evolution, 37(5), 1530–1534. 10.1093/molbev/msaa015

Monadjem, A., Montauban, C., Webala, P., Laverty, T., Moise, B. F., Torrent, L., Tanshi, I., Kane, A., Rutrough, A., Waldien, D., & Taylor, P. (2024). African bat database: Curated data of occurrences, distributions and conservation metrics for Sub-Saharan bats. Scientific Data, 11. 10.1038/s41597-024-04170-7

Mühldorfer, K. (2013). Bats and bacterial pathogens: A review. Zoonoses and Public Health, 60(1), 93–103. 10.1111/j.1863-2378.2012.01536.x

Nyongesa, S. K., Maghenda, M., & Siljander, M. (2022). Assessment of Urban Sprawl, Land Use and Land Cover Changes in Voi Town, Kenya Using Remote Sensing and Landscape Metrics. Journal of Geography, Environment and Earth Science International, 50–61. 10.9734/jgeesi/2022/v26i430347

Omwoyo, A. M., Onwonga, R. N., Wasonga, O. V., & Kinyanjui, M. J. (2024). Spatio-temporal patterns of land use and land cover change in Kibwezi West, Eastern Kenya. Discover Soil, 1(1), 21. 10.1007/s44378-024-00021-4

R Core Team (2026). R: A Language and Environment for Statistical Computing. R Foundation for Statistical Computing, Vienna, Austria.

Rulli, M. C., D’Odorico, P., Galli, N., John, R. S., Muylaert, R. L., Santini, M., & Hayman, D. T. S. (2025). Land Use Change and Infectious Disease Emergence. Reviews of Geophysics, 63(2), e2022RG000785. 10.1029/2022RG000785

Shapiro, J., & Barbier, E. (n.d.). First Record of Streblidae, Raymondia alulata Speiser, 1908 (Diptera: Streblidae), in Swaziland and a Review of the Genus Raymondia and Their Hosts in Africa. 10.3161/15081109ACC2016.18.1.015

Szentivanyi, T., McKee, C., Jones, G., & Foster, J. T. (2023). Trends in Bacterial Pathogens of Bats: Global Distribution and Knowledge Gaps. Transboundary and Emerging Diseases, 2023, 9285855. 10.1155/2023/9285855

Tasker, S. (2020). Hemotropic Mycoplasma. In Clinical Small Animal Internal Medicine (pp. 927–930). John Wiley & Sons, Ltd. 10.1002/9781119501237.ch97

Till, W. M., & Evans, G. O. (n.d.). The genus Chelanyssus Zumpt and Till (Acari: Mesostigmata).

Veikkolainen, V., Vesterinen, E. J., Lilley, T. M., & Pulliainen, A. T. (2014). Bats as Reservoir Hosts of Human Bacterial Pathogen, Bartonella mayotimonensis. Emerging Infectious Diseases, 20(6), 960–967. 10.3201/eid2006.130956

Voigt CC, Phelps KL, Aguirre LF, Corrie Schoeman M, Vanitharani J, Zubaid A. Bats and Buildings: The Conservation of Synanthropic Bats. Bats in the Anthropocene: Conservation of Bats in a Changing World. 2015 Aug 29;427–62. doi:10.1007/978-3-319-25220-9_14 PubMed PMID: null; PubMed Central PMCID: PMC7123121.

Volokhov, D. V., Norris, T., Rios, C., Davidson, M. K., Messick, J. B., Gulland, F. M., & Chizhikov, V. E. (2011). Novel hemotrophic mycoplasma identified in naturally infected California sea lions (*Zalophus californianus*). Veterinary Microbiology, 149(1), 262–268. 10.1016/j.vetmic.2010.10.026

Wang, R., Li, Z.-M., Peng, Q.-M., Gu, X.-L., Zhou, C.-M., Xiao, X., Han, H.-J., & Yu, X.-J. (2023). High prevalence and genetic diversity of hemoplasmas in bats and bat ectoparasites from China. One Health, 16, 100498. 10.1016/j.onehlt.2023.100498

Willi, B., Boretti, F. S., Meli, M. L., Bernasconi, M. V., Casati, S., Hegglin, D., Puorger, M., Neimark, H., Cattori, V., Wengi, N., Reusch, C. E., Lutz, H., & Hofmann-Lehmann, R. (2007). Real-Time PCR Investigation of Potential Vectors, Reservoirs, and Shedding Patterns of Feline Hemotropic Mycoplasmas. Applied and Environmental Microbiology, 73(12), 3798–3802. 10.1128/AEM.02977-06

Windsor, J. J. (2001). Cat-scratch disease: Epidemiology, aetiology and treatment. British Journal of Biomedical Science, 58(2), 101–110.

Zellner, P. N., Njoroge, E., Lynch, A., & Brown, L. D. (2026). Physiology of the cat flea (Ctenocephalides felis) gut: Serine protease activity, midgut pH, and intestinal barrier integrity. Journal of Medical Entomology, 63(2). 10.1093/jme/tjag037

