## Supplemental Figures for "*Bartonella* and hemotropic *Mycoplasma* species in synanthropic bats in Kenya"

**Supplemental Material**

Table S1. Model results investigating the drivers of hemoplasma and Bartonella spp. infection in bats. Odds ratios, 95% confidence intervals, and P-values are provided for fixed effects.

| **Fixed Effect** | ***Bartonella* spp. infection** | | | **Hemoplasma infection** | | |
|---|---|---|---|---|---|---|
| | OR | 95% CI | P | OR | 95% CI | P |
| ***Mops condylurus*** | | | | | | |
| Ectoparasite load | 1.15 | 0.93–1.42 | 0.19 | 1.07 | 0.83–1.38 | 0.58 |
| Sex (male) | 1.17 | 0.72–1.92 | 0.53 | 0.94 | 0.55–1.62 | 0.83 |
| Age class (juvenile) | 0.91 | 0.41–2.03 | 0.81 | 0.32 | 0.09–1.12 | 0.08 |
| Year (2024) | 1.20 | 0.73–1.97 | 0.46 | 0.72 | 0.40–1.29 | 0.27 |
| ***Mops pumilus*** | | | | | | |
| Ectoparasite load | 0.53 | 0.22–1.30 | 0.17 | 0.96 | 0.62–0.60 | 0.86 |
| Sex (male) | 1.22 | 0.42–3.53 | 0.72 | 0.82 | 0.42–1.61 | 0.56 |
| Age class (juvenile) | 1.20 | 0.31–4.60 | 0.80 | 0.74 | 0.30–1.82 | 0.51 |
| Year (2024) | 0.36 | 0.04–3.25 | 0.36 | 0.85 | 2.11–3.43 | 0.82 |
| ***Cardioderma cor*** | | | | | | |
| Ectoparasite load | **2.93** | **1.60–6.23** | **<0.01** | 0.83 | 0.40–1.51 | 0.57 |
| Sex (male) | **0.15** | **0.02–0.79** | **<0.05** | 1.53 | 0.39–5.58 | 0.53 |
| Age class (juvenile) | <0.01 | 0.00–Inf. | 0.99 | 2.44 | 0.40-13.67 | 0.31 |
| Year (2024) | 1.52 | 0.31–7.82 | 0.60 | 2.19 | 0.45–9.77 | 0.31 |

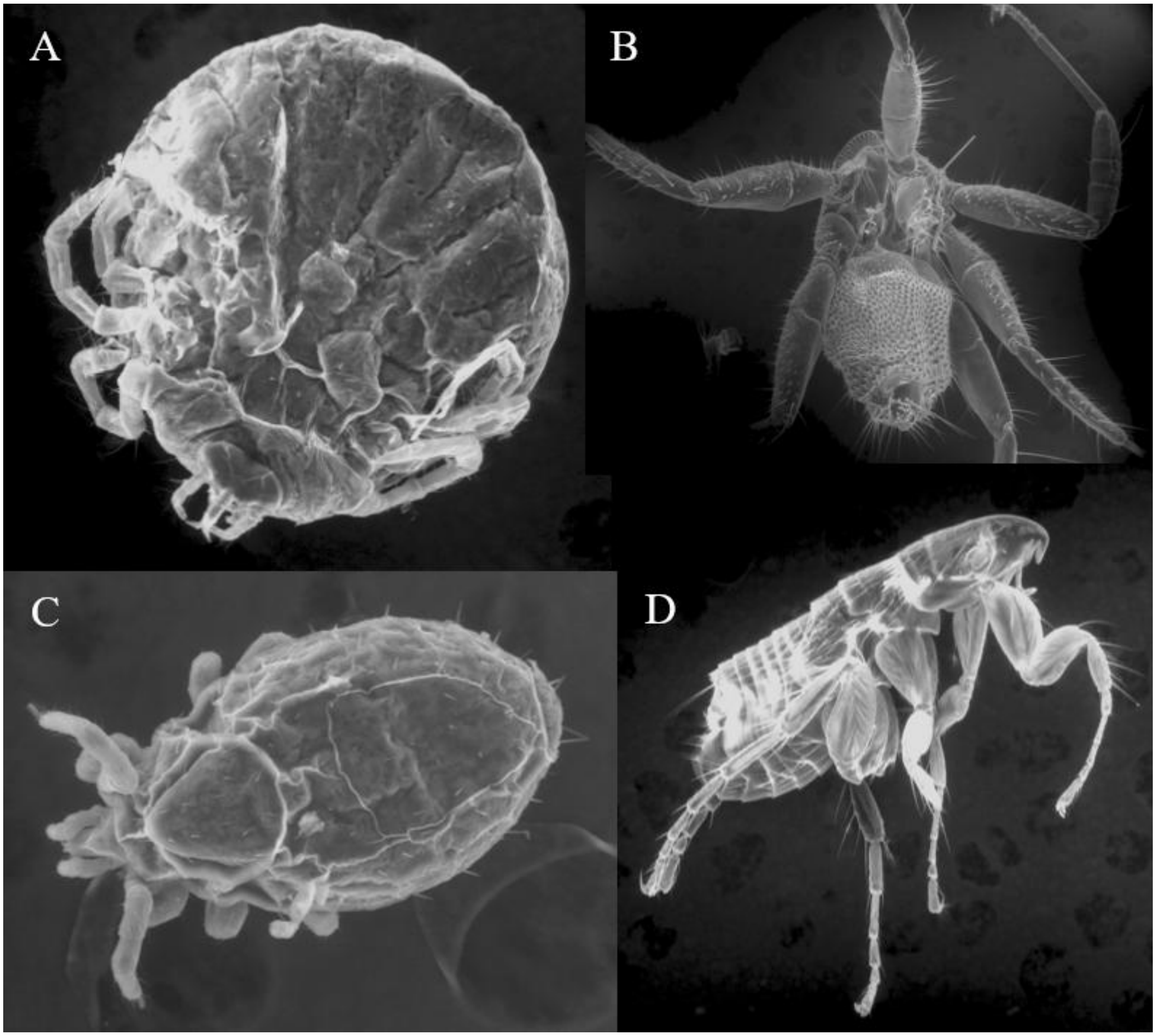


Figure S1. Bat ectoparsites in Taita-Taveta County, Kenya. A. Argasid tick (*Argas boueti)* collected from *Cardioderma cor*; B. Nycteribiid fly (*Raymondia planiceps*) collected from *Cardioderma cor*; C. Macronyssid mite (*Chelanyssus aethiopicus*) collected from *Mops pumilus*; D. Ischnopsyllid flea (*Lagaropsylla idea*) collected from a *Mops condylurus*.
